# Targeting aberrant DNA methylation in mesenchymal stromal cells as a treatment for myeloma bone disease

**DOI:** 10.1101/767897

**Authors:** Antonio Garcia-Gomez, Tianlu Li, Javier Rodríguez-Ubreva, Laura Ciudad, Francesc Català-Moll, Montserrat Martín-Sánchez, Laura San-Segundo, Xabier Morales, Carlos Ortiz de Solórzano, Julen Oyarzabal, Edurne San José-Enériz, Xabier Agirre, Felipe Prosper, Mercedes Garayoa, Esteban Ballestar

## Abstract

Multiple myeloma (MM) progression and myeloma-associated bone disease (MBD) are highly dependent on the bone marrow (BM) microenvironment, in particular on mesenchymal stromal cells (MSCs). MSCs from MM patients exhibit an abnormal transcriptional profile, suggesting that epigenetic alterations could be governing the tumor-promoting functions of MSCs and their prolonged osteoblast (OB) suppression in MM. In this study, we analyzed the DNA methylome of BM-derived MSCs from patients with monoclonal gammopathy of undetermined significance, smoldering myeloma and symptomatic MM at diagnosis in comparison with their normal counterparts. DNA methylation alterations were found at each of the myeloma stage in association with deregulated expression levels of Homeobox genes involved in osteogenic differentiation. Moreover, these DNA methylation changes were recapitulated *in vitro* by exposing MSCs from healthy individuals to MM plasma cells. Pharmacological targeting of DNMTs and G9a with the dual inhibitor CM-272, reverted the expression of aberrantly methylated osteogenic regulators and promoted OB differentiation of MSCs from myeloma patients. Most importantly, in a mouse model of bone marrow-disseminated MM, administration of CM-272 prevented tumor-associated bone loss and reduced tumor burden. Our results demonstrated that not only was aberrant DNA methylation a main contributor to bone formation impairment found in MM patients, but also its targeting by CM-272 was able to reverse MM-associated bone loss.

**KEY POINTS:**

- Bone marrow-derived mesenchymal stromal cells (MSCs) from monoclonal gammopathy of undetermined significance, smoldering myeloma and myeloma patients exhibit an aberrant DNA methylome compared to their healthy counterparts.
- These DNA methylation changes are associated with an altered expression of genes of the Homeobox loci that orchestrate osteogenic differentiation of mesenchymal precursors.
- MM plasma cell-exposed healthy MSCs recapitulate the DNA methylation alterations observed in MSCs isolated from myeloma patients.
- Dual targeting of DNMTs and the histone methyltransferase G9a with CM-272 not only controls MM tumor burden but also prevents myeloma-associated bone loss.

## INTRODUCTION

Multiple myeloma (MM) is an incurable hematological malignancy of clonal expansion of plasma cells in the bone marrow (BM) that accounts for 1% of all cancers ^1, 2^. Nearly 90% of myeloma patients suffer from skeletal related events during the course of the disease, including severe bone pain, hypercalcemia, pathological fractures and spinal cord compression ^3^, that not only affect the quality of life but also the overall survival of MM patients ^4^. Myeloma-associated bone disease (MBD) is characterized by an increase in bone-resorptive activity and number of osteoclasts (OCs), as well as impairment of bone-forming activity and differentiation of osteoblasts (OBs), which ultimately lead to the development of osteolytic lesions ^5^.

In most cases, symptomatic myeloma is preceded by sequential asymptomatic stages of monoclonal gammopathy of undetermined significance (MGUS) and smoldering myeloma (SMM), with increasing BM plasmocytosis and monoclonal component as well as augmented risk of progression to active MM ^6 7^. The biological behavior and clinical outcome of MM is partly dependent on genetic and epigenetic abnormalities of tumor subclones that arise from MGUS and SMM stages^8^. However, the clinical stability of MGUS cases, despite displaying shared genetic lesions with MM cells, suggests that the BM microenvironment may critically regulate disease progression ^9 6 10^. In this regard, it has been widely shown that a complex and bidirectional relationship exists between MM cells and the BM niche, which results in oncogenesis support, anaemia, immunosuppression and uncoupling of the bone remodeling process ^11^.

Mesenchymal stromal cells (MSCs) are an essential cell type in the formation and function of the BM microenvironment, being the progenitors of bone-forming OBs, adipocytes and chondroblasts, as well as the haematopoietic-supporting stroma components of the BM ^12^. It is well-documented that BM-derived MSCs from MM patients contribute to MM progression (reviewed in ^11^) and show an impaired ability to differentiate into OBs ^13 14^. Moreover, MM-MSCs are considered inherently abnormal, as their dysfunctionality remains even following *ex vivo* culture in the absence of MM cells^15^. Furthermore, bone lesions persist in many MM patients even after therapeutic remission, suggesting a long-term defect in MSCs that inhibit their ability to properly differentiate into functional OBs ^16^.

Previous studies described that MSCs from MM patients are cytogenetically normal ^17 18^, but show alterations in their transcriptional ^13 19^ and proteomic ^11^ profiles even in the absence of myeloma cell interaction. This suggests that epigenetic mechanisms could be governing the tumor-promoting functions of MSCs and their prolonged OB suppression in MM. In fact, Adamik and colleagues reported an abnormal recruitment of chromatin remodelers in MSCs from myeloma patients, contributing to the transcriptional repression of Runx2, a master regulator of OB differentiation ^20^. Yet, there is a lack of information about DNA methylation-based mechanisms that may contribute to MM progression and subsequent bone defects. DNA methylation is an essential epigenetic modification involving the addition of a methyl group to the 5-carbon of the cytosine ring by a family of DNA methyltransferase (DNMT) enzymes ^21^, which has been described to play a critical role in MSC lineage determination ^22^, as well as in tumor progression and immunosuppression in other cancer types ^23^..

In this study, we aimed to determine the DNA methylome of BM-derived MSCs in the context of MM, since not only would they play an essential role in OB differentiation but would also perpetuate an aberrant functional state of the BM niche and drive disease progression. We identified DNA methylation alterations in MSCs and the ability of MM plasma cells to induce such changes, which result in the dysregulation of critical genes for osteogenesis. Finally, we explored the *in vitro* and *in vivo* preclinical efficacy of CM-272, a dual inhibitor of DNMTs and the histone methyltransferase G9a, which cooperates closely with DNA methylation, in the context of MBD.

## METHODS

### Participants

BM samples were obtained from patients with newly diagnosed MGUS (n = 10), SMM (n = 8), and MM (n = 9), according to the International Myeloma Working Group criteria. BM samples from healthy controls (n=8) were obtained from participants undergoing orthopedic surgery. Each sample was obtained after receiving informed written consent of all participating subjects and following approval from the University Hospital of Salamanca de Salamanca Review Board. Clinical characteristics of MGUS, SMM and MM patients are listed in **Supplementary Table S1**.

### Statistical analysis

DNA methylation and transcriptomic array analyses were performed using the R environment, in which a student t test was used to determine statistically significant differences between groups. Each *in vitro* assay was performed using MSCs from at least three donors. Data are expressed as mean ± SEM. Statistical analyses were carried out with Prism version 6.0 (GraphPad) and were performed using a two-tailed Mann-Whitney *U* test or Student’s *t* test.

Information regarding reagents, cells, methylation/sequencing/bioimaging platforms, molecular biology techniques and *in vivo* studies are described in **Supplementary Methods**.

## RESULTS

### BM-derived MSCs of distinct MM stages exhibit altered DNA methylation profiles

We first obtained genome-wide DNA methylation profiles of BM-derived MSCs isolated at different stages of MM (newly diagnosed MGUS, high-risk SMM and MM) and healthy controls. DNA methylation changes were identified using two different statistical approaches **(Figure 1A)**: i) detection of differentially methylated CpG positions (DMPs) based on differences in DNA methylation means between patient (MGUS, SMM and MM) and healthy MSCs (Δβ ≥ 0.15 and \*\**p* < 0.01) **(Supplementary Table S2)**; and ii) detection of differentially variable CpG positions (DVPs) based on differences in variance of DNA methylation levels (FDR < 0.05 and \**p* < 0.05) between the sample groups **(Supplementary Table S3)**. In regards to DMPs, the largest number of altered CpGs was found in the SMM stage compared to healthy donors **(Supplementary Figure S1A and S1B)**. On the other hand, we observed the highest number of DVPs in MGUS followed by SMM and MM samples **(Supplementary Figure S1C and 1D)**, supporting the notion that these stochastic and heterogeneous DNA methylation patterns are associated with early stages of carcinogenesis, as previously reported ^24 25^. We also observed that the majority of identified DMPs and DVPs are disease stage-specific, although the asymptomatic stages showed a moderate proportion of overlap **(Figure 1A)**.

**Figure 1.**
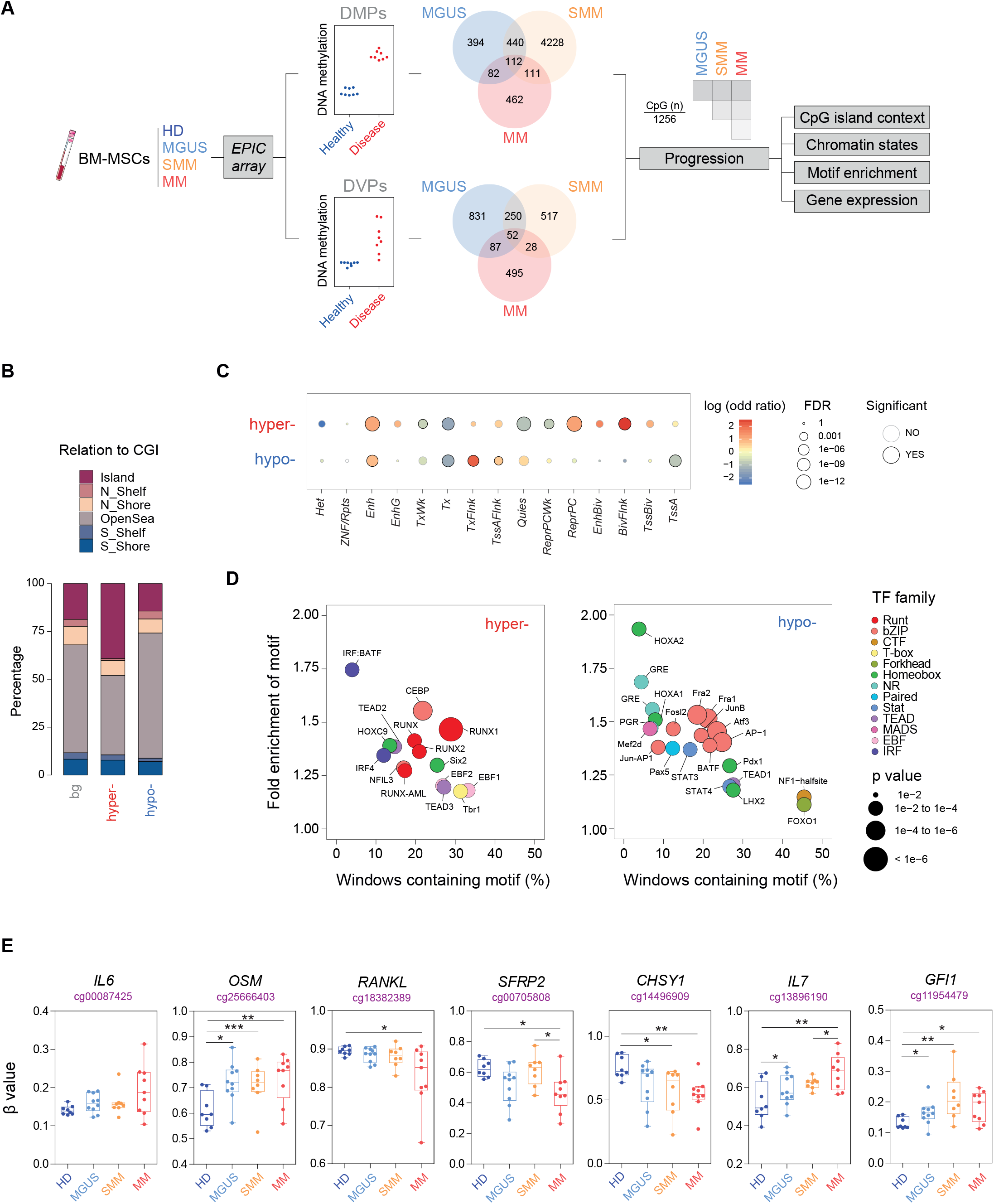
High-throughput stepwise DNA methylation changes in BM-derived MSCs associated to MM progression. A) Workflow depicting the metholodogical approach for selecting DNA methylation changes in BM-derived MSCs from MGUS, SMM and MM patients versus healthy controls. An example of CpG site experiencing increased mean (DMP) or variance (DVP) in the disease versus the control condition is shown. Venn diagrams show the number of DMPs or DVPs resulting from each comparison. B) Distribution of DNA methylation changes in relation to CpG islands (CGI), including shores, shelves and open sea regions for differentially hyper- or hypo-methylated CpG sites. C) Enrichment analysis of differentially hyper- and hypo-methylated CpG sites located in different genomic regions, annotated by 15 chromHMM states. Color scale refers to log odd ratio and circle size refers to *p* value significance. D) Bubble plot representation of HOMER transcription factor (TF) motif enrichment analysis of differentially hyper- and hypo-methylated CpGs in MSCs during MM progression (left and right panel respectively). Colour range depicts different transcription factor families and circle size refers to *p* value significance. E) Box-plots showing β-values obtained from the EPIC array in MSCs from healthy donors and MGUS, SMM and MM patients of relevant genes involved in the pathogenesis of MM and associated bone disease.

Given that myeloma is a multi-stage disease, we then analyzed the accumulative changes of DNA methylation associated to MM progression by selecting DMPs and DVPs that were found either only in MM stage, shared by SMM and MM and in all three stages (574 hyper- and 682 hypo-methylated CpGs) **(Figure 1A) (Supplementary Table S4)**.

Analyzing the distribution of MM progression-associated CpGs in relation to CpG islands (CGI), we observed a significant enrichment of CpG islands in the hypermethylated CpG set **(Figure 1B)**. Utilizing publicly available chromatin state maps of BM-derived MSCs from healthy individuals ^26^, we found a significant enrichment of both hyper- and hypo-methylated CpG sites that correspond to enhancers **(Figure 1C)**. In addition, we observed an enrichment in flanking transcription start sites (TSS) in the hypomethylated set, and bivalent TSS and Polycomb Group (PcG) genes in hypermethylated CpGs **(Figure 1C)**.

To determine whether these MM progression-associated loci shared any common DNA elements, we performed a search for enriched transcription factor (TF)-binding sites in these regions using the HOMER algorithm ^27^. We observed a significant over-representation of binding sites for Runt and Tead family in differentially hyper-methylated regions associated to MM progression (*\*\*p*<0.01; **Figure 1D**). These results suggest that key transcription factors involved in the upregulation of osteogenic genes, such as RUNX2 ^28^ or TEAD2 ^29^, might participate in aberrant DNA hypermethylation. Since DNA methylation has been originally linked to transcriptional repression, these results suggested that the hypermethylation of these regions could compromise the ability of MSCs to undergo proper differentiation. On the other hand, CpG sites that experience aberrant DNA hypomethylation were highly enriched in binding motifs of the bZip and Homeobox families (*\*\*p*<0.01; **Figure 1D**). In this respect, the loss of DNA methylation could be selectively driving the occupancy of TF that have been reported as negative regulators of OB differentiation such as HOXA2 ^30^ and ATF3 ^31^. In addition, we observed transcriptional deregulation of some members of these TF families using expression array data from BM-derived MSCs of healthy controls, MGUS, SMM and MM patients. Some of these TFs were specifically downregulated in MSCs of active myeloma (*RUNX2* and *TEAD2*), whereas others were already downregulated in precursor myeloma stages (*HOXC9* and *CEBPD*) **(Supplementary Figure S1E)**. In all, these findings suggested that MM progression-associated DNA methylation changes in MSCs were mediated by the sequential activity of specific TF families, which were also functionally deregulated in MM ^32^. Furthermore, other genes that play important roles in the pathophysiology of MM (such as the cytokines IL6 and OSM) and associated MBD (secreted factors such as RANKL, SFRP2, IL7, CHSY1 and the transcriptional repressor GFI1) were also found to alter their DNA methylation levels **(Figure 1E)**.

### Aberrant DNA methylation is associated with differential Homeobox gene expression in MSCs at different MM stages

To further investigate the relationship between differential DNA methylation and gene expression, we mapped the DMPs and DVPs **(Supplementary Table S2 and S3)** to the most proximal gene. Using expression array data from BM-derived MSCs of healthy controls, MGUS, SMM and MM patients **(Supplementary Table S5)**, differential expression of DMP- and DVP-associated genes were identified using a cutoff of \**p* < 0.05 comparing MGUS/SMM/MM to healthy controls **(Figure 2A) (Supplementary Table S6).** Gene ontology (GO) analysis revealed that the genes displaying both differential methylation and expression were enriched in functional categories important in cell fate commitment and bone phenotype **(Figure 2B)**. The most enriched functional category corresponded to genes from the Homeobox family. Within the Homeobox family, we found the subset of *Hox* genes that encodes a large family of highly conserved TFs responsible for driving the correct differentiation of MSCs ^33^, namely genes belonging to the HOXA-to-D clusters. Furthermore, we observed a significant enrichment in genes reported to be downregulated in MM-MSCs (Figure 2B) ^13^. Integration of methylation and gene expression data corresponding to the Homeobox and bone formation-related genes revealed that both DNA hyper- and hypo-methylation events were associated with both gene downregulation and upregulation **(Figure 2C)**. Specifically, hypermethylated genes that showed a reduced expression in patient MSCs include positive regulators of OB differentiation such as *RUNX2* or *NRP2* ^34^ **(Figure 2C)** In contrast, negative regulators of osteogenesis such as *SFRP2* ^35^ *or NFATC2* ^36^ were hypomethylated and consequently upregulated in patient MSCs. In all, these factors could potentially contribute to impaired osteoblastogenesis associated to bone disease in MM **and this is summarized in Supplementary Table S7)**.

**Figure 2.**
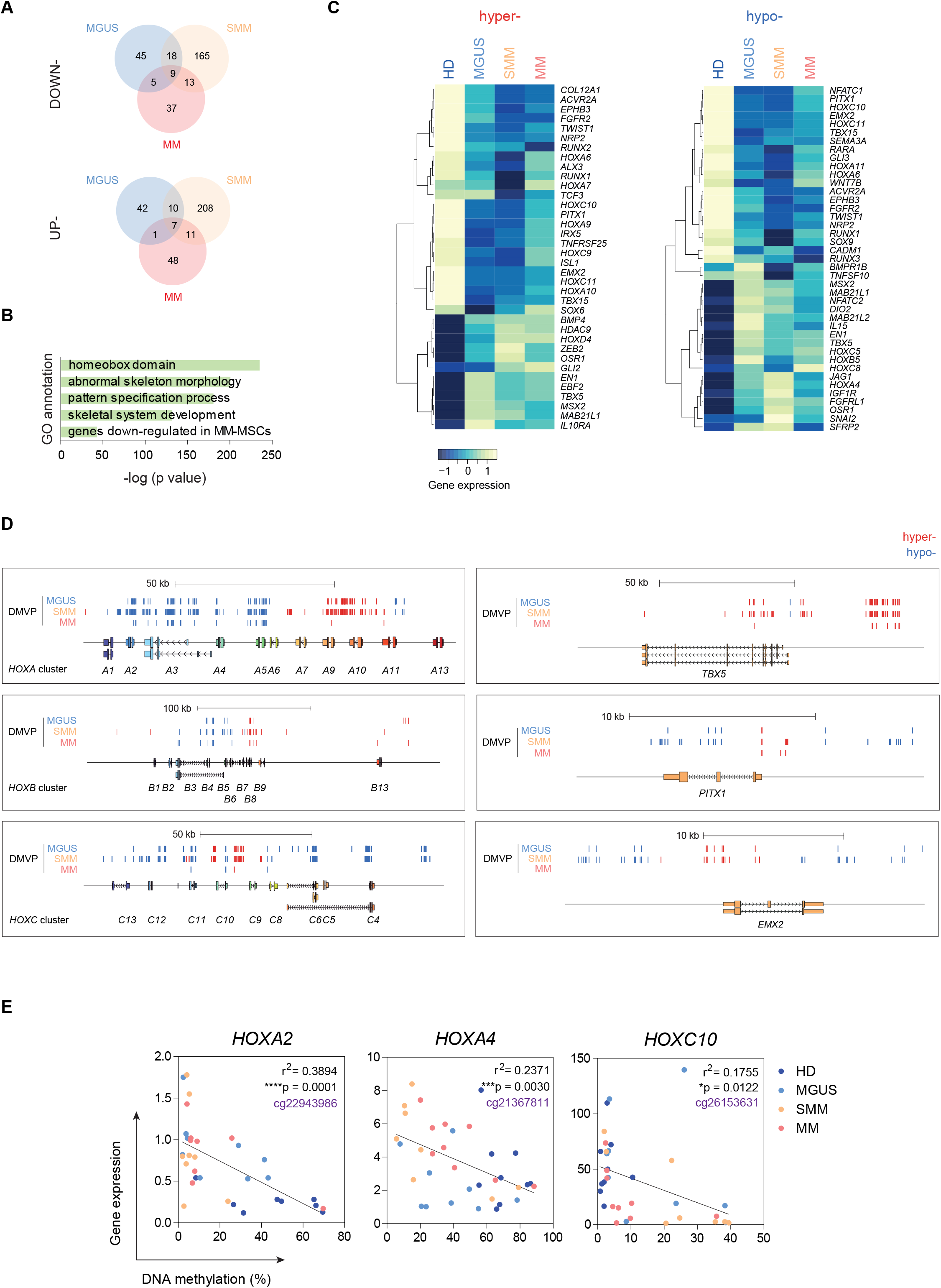
DNA methylation changes associate with differential gene expression of Homeobox genes in MSCs from MGUS, SMM and MM patients. A) Venn diagrams showing differentially methylated or down-regulated (upper) and up-regulated (lower panel) genes when comparing MGUS/SMM/MM samples with healthy individuals. B) Gene ontology (GO) enrichment analysis of CpG sites undergoing DNA methylation and gene expression changes in MSCs of the disease condition. C) Heatmaps showing Homeobox and other OB-related genes associated to differentially hyper-methylated (left) or hypo-methylated (right) CpG sites. D) Scheme depicting differentially methylated CpG sites located in HOX clusters (*HOX-A*, *-B* and *-C*) and other Homeobox genes (*TBX5, PITX1* and *EMX2*). Blue lines indicate hypo-methylated CpGs and red lines indicate hyper-methylated CpGs associated to MGUS, SMM and MM condition. D) Correlation between DNA methylation and gene expression at selected HOX loci. DNA methylation levels of differentially methylated CpG sites were validated by pyrosequencing and gene expression levels of associated HOX genes were validated by qRT-PCR. Pearson correlation coefficient and r2 value were represented. Statistically significant tests are represented as *, *p* < 0.05; ** *p* < 0.01; *** *p* < 0.005.

Upon a closer inspection of several Homeobox-associated genomic regions, we observed a negative correlation between DNA methylation of promoters and gene expression. Specifically, HOXA gene cluster showed aberrant hypo-methylation at the *HOXA4* promoter, and its gene expression was upregulated at different disease stages. Conversely, gene promoters of *HOX-A6, -A7, -A9, -A10* and *-A11* displayed hypermethylation and these genes were downregulated in MGUS/SMM/MM **(Figure 2D) (Supplementary Figure S2)**. A similar pattern of inverse association between methylation and expression was observed in the HOXB and HOXC gene cluster, where *HOXB5, -B6, -C5* and *-C8* were aberrantly hypomethylated and upregulated, whereas *HOXC9, -C10* and *-C11* were hypermethylated and downregulated in patients **(Figure 2D) (Supplementary Figure S2)**. Other Homeobox genes such as *TBX5*, *PITX1* or *EMX2* were also reported as regulators of bone formation ^37 38 39^ and showed an association between DNA methylation at gene promoter and gene expression **(Figure 2D) (Supplementary Figure S2)**.

We then validated the aforementioned DNA methylation and gene expression changes in an independent cohort of BM-derived MSCs from different MM disease stages by pyrosequencing and real-time quantitative PCR assays. Among the differentially methylated genes of the Homeobox family, we selected *HOXA2, -A4* and -*C10* on the basis of their reported role in MSC pluripotency ^40^. In all cases, we observed that DNA methylation negatively correlated with gene expression **(Figure 2E)**.

### Upon interaction with MM plasma cells, healthy MSCs change their DNA methylation profile to one that partially resembles MM patients upon interaction with MM plasma cells

To address the potential contribution of MM cells in mediating aberrant DNA methylation changes in MSCs, we evaluated whether the epigenetic changes observed in MM-MSCs could be mimicked *in vitro* by direct contact of healthy MSCs with MM cells. Thus, we co-cultured BM-derived MSCs from healthy donors with the human MM cell line MM.1S for two weeks Subsequently, MSCs were sorted by CD13+ expression and subjected to DNA methylation analysis **(Figure 3A)**.

**Figure 3.**
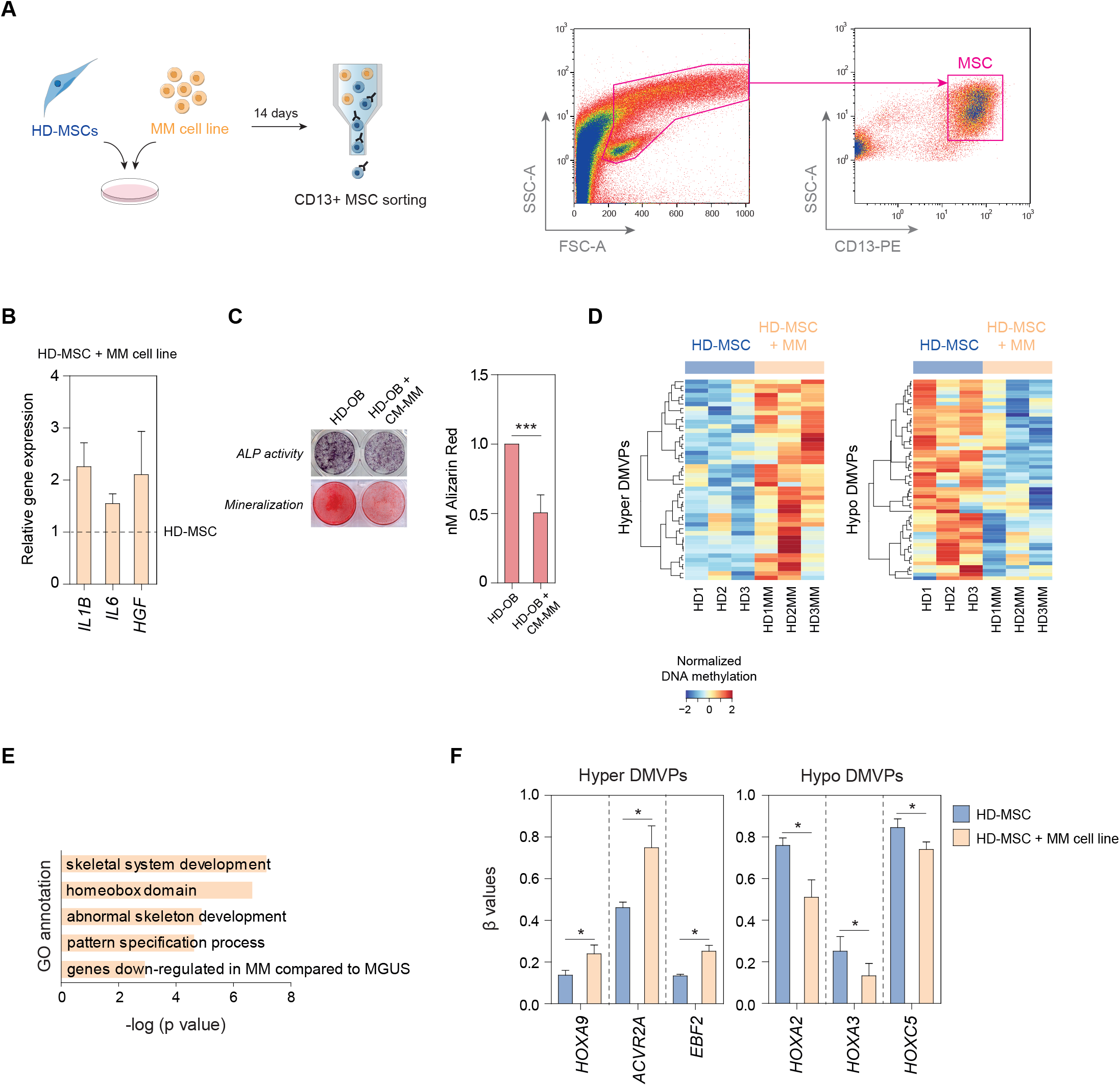
MSCs from healthy donors recapitulate DNA methylation changes observed in MSCs from MM patients upon interacting with MM plasma cells. A) Sorting-strategy for selecting CD13+ MSCs after 14 days of co-culture with the MM.1S cell line. B) Total RNA from the hMSC-TERT cell line was isolated after 14 days in mono-culture or co-culture with the MM.1S cell line, and expression levels of *IL1B*, *IL6, HGF* and *NRP3* were evaluated using qRT-PCR. Data are represented as the mean ± SEM from three different experiments. C) ALP activity and matrix mineralization were assessed in differentiated OBs from HD-MSCs in the presence or absence of conditioned media from the MM.1S cell line. D) Heatmap showing differentially methylated CpG sites (*p < 0.05) in sorted HD-MSCs in mono-culture or co-cultured with the MM.1S cell line for 14 days. E) GO enrichment analysis of CpG sites undergoing DNA methylation in HD-MSCs co-cultured with the MM.1S cell line. F) Bar-plots showing β-values obtained from the DNA methylation array.

Under these conditions, MM.1S cells were able to induce the expression of genes known to be upregulated in MM-MSCs (*IL1B, IL6* and *HGF*) in HD-MSCs compared with mono-cultured HD-MSCs **(Figure 3B)**. Additionally, we validated the inhibitory effect of MM cells in MSC-to-OB differentiation and observed a decrease in both ALP activity and OB mineralization in OBs differentiated in the presence of conditioned media from the MM.1S cell line as compared to OBs differentiated alone.

We then investigated the DNA methylation profiling of MSCs from healthy donors generated upon interaction with MM cells. We observed that 142 CpGs that change their methylation levels upon co-culture with MM.1S cells were shared with aberrant DNA profiles found in MSCs isolated from MGUS/SMM/MM patients **(Figure 3D; Supplementary Table S8)**. Although this accounted for a small percentage of DMPs identified in the *in vitro* study, GO analyses revealed an enrichment in Homeobox genes and categories related with bone formation, similar to what was observed in primary patient MSCs **(Figure 3E)**. Specifically, we found that healthy MSCs exposed to MM cells underwent gains (*HOXA9, ACVR2A, EBF2*) and losses (*HOXA2, HOXA3, HOXC5*) of DNA methylation in the direction of those observed for MSCs from myeloma patients **(Figure 3F)**. Altogether, these results support the notion that MM cells not only are capable of inducing changes in the global methylome of MSCs but also have a significant impact at specific osteogenic loci.

### Dual targeting of DNMTs and G9a restores Homeobox gene expression *in vitro* and promotes osteogenic differentiation of mesenchymal precursors

Gene expression analysis of DNMTs in MSCs from HD and MM patients co-cultured with MM cells obtained from a previous study ^41^ showed an aberrant upregulation of the DNA methyltransferase DNMT1 **(Figure 4A)**. DNMT1 interacts with the methyltransferase G9a to coordinate DNA and H3K9 methylation during cell replication ^42^ promoting transcriptional silencing of target genes. Moreover, G9a can suppress transcription by inducing DNA methylation in addition to its activity as a chromatin remodeler ^43^. In this regard, we hypothesize that the dual inhibition of DNMT1 and G9a could reactivate hypermethylated and silenced genes of MSCs from MM patients preserving their osteogenic potential and therefore preventing myeloma-associated bone loss. Thus, we utilized a dual inhibitor of DNMTs and G9a, termed CM-272, which has been previously described to have a potent therapeutic response, both *in vitro* and *in vivo,* in other neoplasias ^44 45 46 47^.

**Figure 4.**
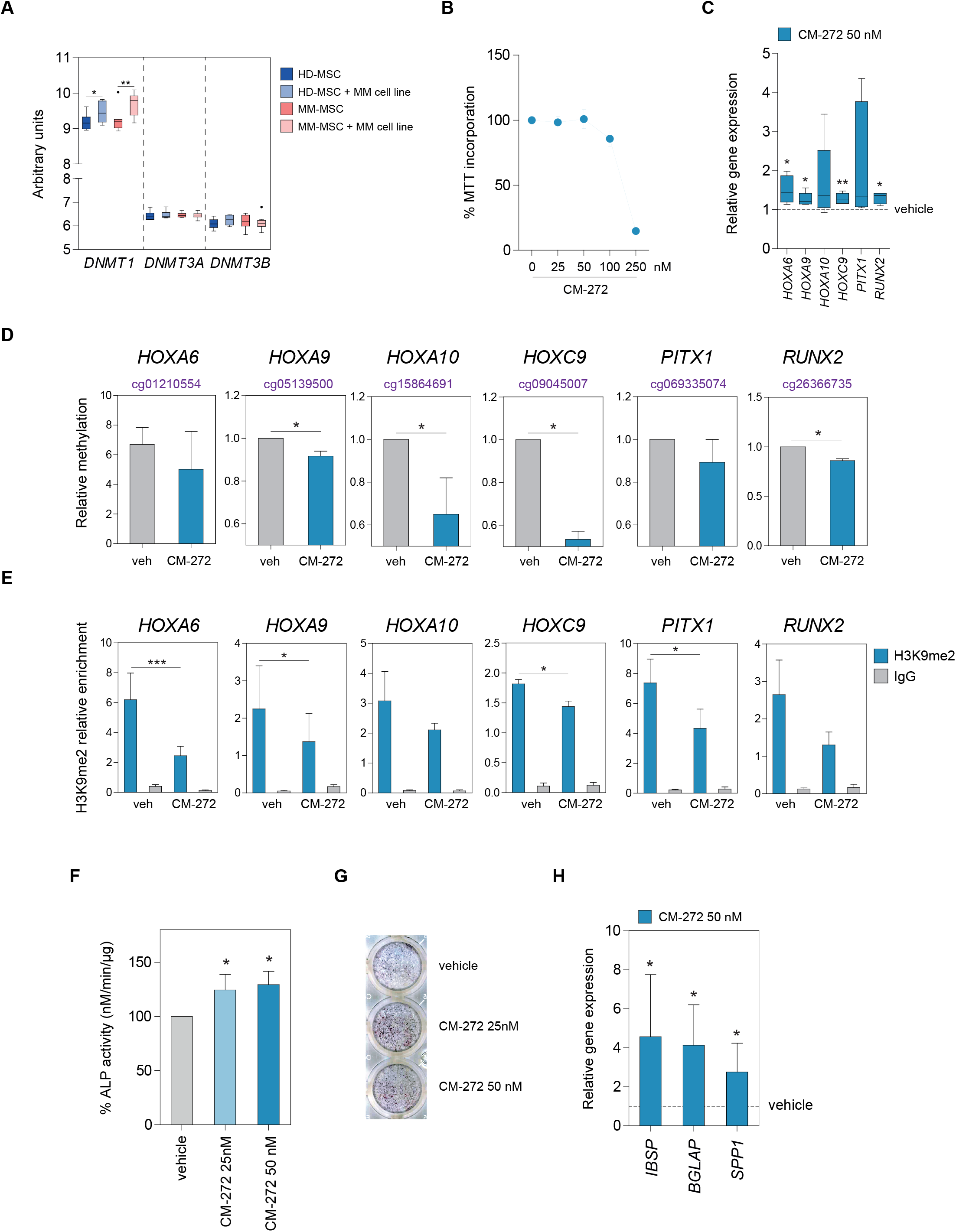
CM-272 treatment reactivates Homeobox gene expression and promotes the osteogenic differentiation of MSCs from MM patients. A) Expression of DNA methyltransferases in MSCs co-cultured for 24 hours with the MM.1S cell line relative to that in mono-culture (from HD and myeloma patients) as assessed by the GeneChip Human Gene 1.0 ST Array. B) MSCs from MM patients were treated with the indicated doses of CM-272 for 72 hours and subjected to MTT assay for viability. C) Real-time RT-PCR was performed to determine the expression of hypermethylated and silenced Homeobox genes (*HOX-A6, -A9, -A10, -C9, PITX1, RUNX2*) in MM-MSCs treated with vehicle or CM-272 for 7 days. D) DNA methylation analysis by pyrosequencing of selected CpGs located at the promoter regions of Homeobox genes in MM-MSCs treated with vehicle or CM-272 for 7 days. E) ChIP assays showing the H3K9me2 enrichment at the promoter regions of Homeobox genes in MM-MSCs treated with vehicle or CM-272 for 7 days. IgG was used as a negative control. Data are shown as relative enrichment of the bound fraction with respect to the input DNA. ALP activity was assessed by F) p-NPP hydrolysis and G) NBT-BCIP staining in MM-MSCs cultured in osteogenic media in the presence or absence of CM-272. H) Expression of osteoblastogenic markers (bone sialoprotein, osteocalcin and osteopontin) was checked by qRT-PCR in MM-MSCs cultured in osteogenic media in the presence or absence of CM-272. Data are shown as mean values of three independent experiments ± SEM. Statistically significant tests are represented as *, *p* < 0.05; ** *p* < 0.01; *** *p* < 0.005 between vehicle and CM-272 condition.

We first checked the effect of CM-272 on the cell viability of mesenchymal progenitors and we selected a dose (50 nM) with no significant toxicity in order to perform further experiments **(Figure 4B)**. CM-272 treatment was able to restore the expression of Homeobox genes (*HOX-A6, -A9, -A10,-C9, PITX1* and *RUNX2*) that were epigenetically repressed in MSCs from MM patients **(Figure 4C)**. Mechanistically, we observed a loss of DNA methylation in the promoter region of the majority of the aforementioned genes after CM-272 treatment in MM-MSCs **(Figure 4D)**. We then checked the levels of the inactive chromatin mark H3K9me2, a hallmark of methyltransferase G9a activity, at these gene promoters upon CM-272 treatment. ChIP-qPCR analysis showed a decrease in H3K9me2 levels at the promoter regions of Homeobox genes after CM-272 treatment **(Figure 4E)**. Taken together, our results suggest that CM-272 acts *in vitro* by inhibition of both DNMT and G9a methyltransferase activity.

Next, we addressed whether targeting DNMT and G9a may have a role in regulating osteogenic differentiation. For this purpose, we cultured MSCs from myeloma patients in osteogenic media to obtain differentiated OBs in the presence or absence of CM-272. As observed in **Figure 4F** **and** **G**, CM-272 was able to increase ALP activity in early-stage OBs. Furthermore, CM-272 treatment was able to upregulate the relative expression of several late bone formation markers (namely, bone siaploprotein, osteopontin, and osteocalcin) in MSCs from myeloma patients **(Figure 4H)**.

### CM-272 not only controls tumor burden but also prevents the myeloma-associated bone loss

To test the effect of CM-272 in the context of MBD, we used an established murine model of bone marrow-disseminated myeloma. After equivalent engraftment of myeloma cells (RPMI8226-luc) was verified by bioluminescence measurement, mice were treated for 4 weeks with CM-272 as described in Methods. Compared with the vehicle control group, CM-272 controlled tumor progression as measured by bioluminescence **(Figure 5A)** or by serum levels of hIgλ secreted by MM cells **(Figure 5B)**. Representative microCT images at the metaphyses of distal femurs showed tumor-associated bone loss in vehicle-treated mice, in contrast with trabecular structures observed in CM-272-treated animals **(Figure 5C)**. The 3D reconstruction images of distal femurs revealed a marked bone loss evidenced by a thin trabecular network (in red) but also by loss of cortical bone (in grey) in vehicle-treated mice **(Figure 5D)**. By contrast, CM-272-treated mice presented a gain in both trabecular and cortical bone **(Figure 5D)**. This was also reflected by bone morphometric parameters which resulted in increased trabecular bone volume, occupancy and connectivity and reduced trabecular separation in CM-272–treated animals, as compared with vehicle control **(Figure 5E)**. Finally, these findings correlated with a significant increase in serum levels of the bone formation marker P1NP analyzed after CM-272 treatment compared to untreated control **(Figure 5F)**. In summary, these data demonstrate that CM-272 exerts *in vivo* anti-myeloma activity along with bone anabolic effects in human MM bearing-mice.

**Figure 5.**
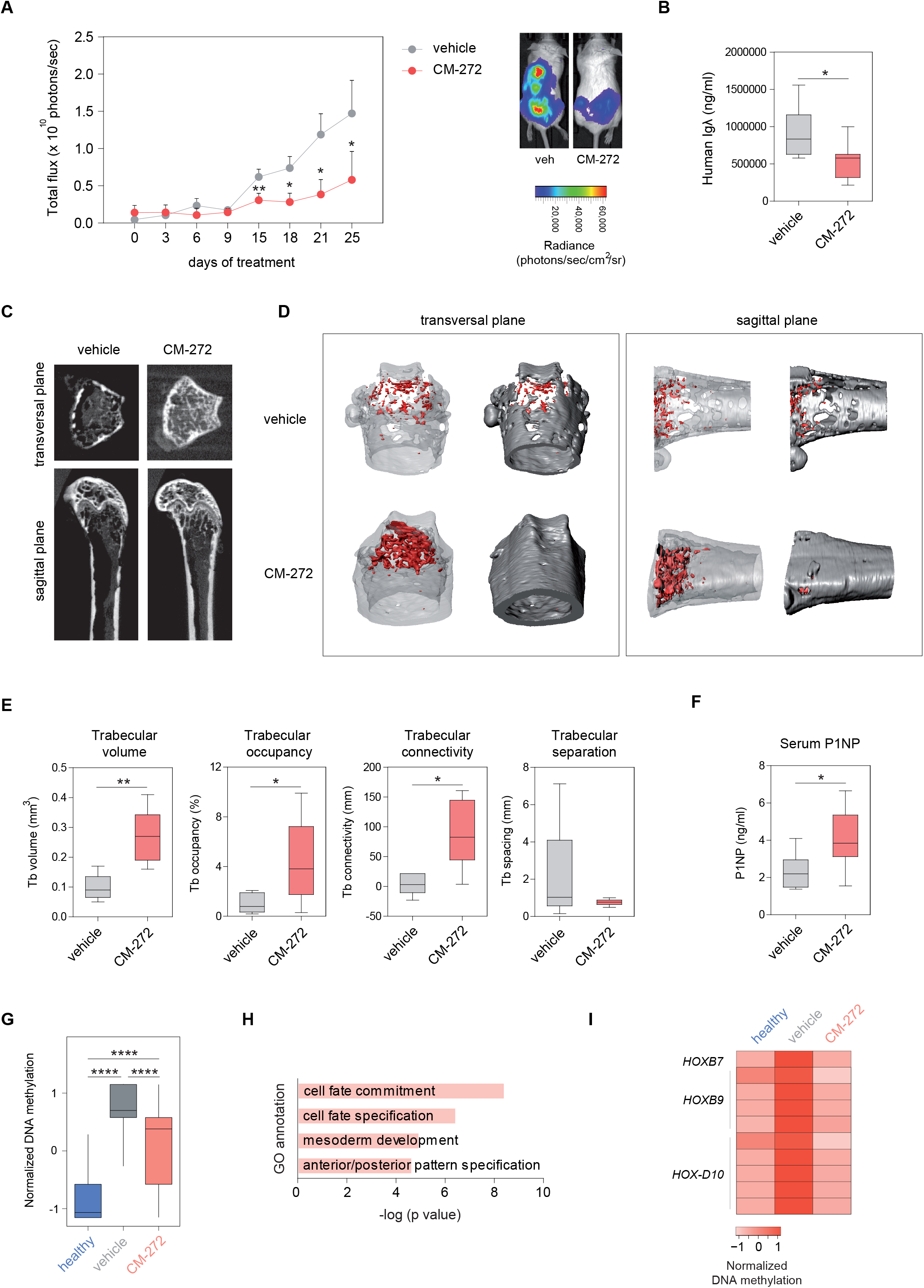
CM-272 prevents tumor-associated bone loss besides reducing multiple myeloma tumor burden. A) RPMI8226-luc cells (8 × 10^6^) were intravenously injected into NSG mice. After 4 weeks, mice were randomized into 2 groups [receiving vehicle and CM-272; *n* = 6/group], and treated for additional 4 weeks with dosing and regimen schedules as specified in Supplementary Methods. Tumor dissemination was checked by A) bioluminescence measure and B) serum levels of human Igλ secreted by RPMI8226-luc cells at specified time points. C) Representative microCT cross-sections at the metaphyses of distal femurs in vehicle and CM-272-treated mice in transversal (upper) and sagittal (down) planes. D) Transversal (left) and sagittal (right) planes of corresponding 3D renderings from microCT images at distal femurs (trabecular bone in red, cortical bone in grey). E) Trabecular bone morphometric parameters from micro-CT images were quantitated for trabecular bone volume, occupancy, connectivity and separation. F) Serum levels of the bone formation marker P1NP were quantified by ELISA. Graphs represent mean values ± SEM. *, *p* < 0.05; ** *p* < 0.01 versus the vehicle control group. G) Box-plots showing DNA methylation levels of pooled MSCs obtained from healthy, vehicle- and CM-272-treated animals corresponding to hypermethylated CpGs between healthy and tumor-bearing animals. H) GO enrichment analysis of CpG sites undergoing DNA hypermethylation changes in vehicle-treated MSCs versus MSCs from healthy mice. I) Heatmap showing normalized DNA methylation levels of individual CpGs at selected Homeobox loci among animal groups. Data pooled from mice (n=7) for each group with sufficient RRBS coverage (≥5 valid sequencing reads per CpG).

To further examine the *in vivo* effect of CM-272 on DNA methylation of myeloma-associated MSCs, we performed reduced representation bisulfite sequencing (RRBS) analysis of MSCs isolated from vehicle- and CM-272-treated myeloma-bearing mice using healthy mice as controls. First, we observed significant alterations in the DNA methylome of myeloma bearing mice compared to healthy mice **(Supplementary Table S9)**. We then compared the methylation status of hypermethylated CpGs in mice treated with CM-272 and observed a partial reversal of the aberrant hypermethylation **(Figure 5G)**. These DNA methylation changes occurred at genomic loci enriched for genes involved in cell commitment and differentiation, such as Homeobox genes **(Figure 5H)**. Specifically, we were able to identify CpGs that experienced a gain in DNA methylation in vehicle-treated MSCs compared to healthy controls at the same genomic loci previously identified in human MM-MSCs (*HOX-B7, -B9, -D10*). Importantly, CM-272 treatment was able to restore the DNA methylation levels at these loci to resemble that of healthy mice, which was concomitant with the reduced tumor burden as well as bone loss recovery observed in these mice. **(Figure 5I)**.

## DISCUSSION

The pathogenic transition from premalignant stages to active MM is complex and not well understood. One example of this complexity is that although all MM cases emerge from pre-existing asymptomatic MGUS/SMM stage, not all MGUS progress into MM and can exist as a stable and independent disease. Nevertheless, despite being an asymptomatic stage, transformed plasma cells in MGUS present cytogenetic alterations similar to that of myeloma plasma cells, as well as significant abnormalities in bone remodeling ^48 49^. This indicates that both genetic and microenvironmental alterations exist from early stages of the disease. In our study, we show that epigenetic alterations in MSCs already occur in early asymptomatic stages of both MGUS and SMM, and although many alterations are shared between all stages, the majority of DNA methylation changes are specific to each stage. These results are in accordance with previous studies that indicate the existence of stage-specific epigenetic alterations during MM progression in malignant plasma cells ^50 51^. This phenomenon could be explained by the expansion of sub-populations of MSCs during MM disease progression which may favor tumor development and drug resistance, similarly to what was observed to occur in MM cells ^52 53^.

Deregulation of methylome in MM-MSCs mediates transcriptional and phenotypical alterations. Interestingly, many genes of the Homeobox family displayed both epigenetic and transcriptional dysregulation in patient MSCs, and these changes were observed in earlier stages of disease. In this regard, members of the HOX family have been recently described to be key drivers of OB differentiation, in which their expression is fine-tuned by demethylation of their promoters during the osteogenic process ^54^. Furthermore, we observed that healthy MSCs exposed to MM cells, similar to what was observed in patient MSCs, not only displayed an altered methylome, but also showed impaired MSC-to-OB differentiation, as previously described ^20^. Hence, our results suggest that the impairment of osteogenesis in all stages of MM arise from early transcriptional deregulation of Homeobox genes, in which altered DNA methylation may be the primary mediator in this process.

Although the biology of MBD is relatively well-described, there is still a lack of pharmacological treatments to improve bone loss. Clinically approved bone-modifying agents for the treatment of MBD include bisphosphonates ^55^, that inhibit bone resorption by suppressing OC activity, and denosumab ^56^, a monoclonal antibody against the osteoclastogenic cytokine RANKL. However, these drugs only target the OC compartment, and bone disease still persists due to the absence of bone formation. Thus, therapeutic agents targeting OBs are needed. In this study, we demonstrate a new strategy for treating MBD by targeting aberrant DNA methylation in MSCs. Firstly, we observed that treatment with CM-272 was able to reverse the transcriptional and epigenetic alterations in MSCs from myeloma patients. Additionally, this agent promotes the ability of MSCs to differentiate into OBs. These *in vitro* effects on bone were mirrored in a mouse model of disseminated MM. Of note, CM-272 treatment not only prevents bone loss by bone-anabolic effects but also show anti-myeloma activity. This is in line with previous reports showing that DNMTs are targets for the treatment of MM ^57 58 59^ and also for improving the osteogenic differentiation ability of MSCs ^60^. Additionally, we cannot discard the possibility that the observed effects on tumor growth inhibition may be a consequence of the impairment of the cross-talk between MSCs and MM cells. Moreover, the dual targeting effects of CM-272 also inhibit the dimethylation of H3K9, which has been described to be crucial in the establishment of DNA methylation ^42 43^. It is therefore rational to envision that the bone anabolic effects mediated by CM-272, both *in vitro* and *in vivo*, involves the reversion of aberrant hypermethylation at Homeobox loci in the MSC population.

In summary, our findings highlight the existence of aberrant DNA methylation patterns in the BM-derived MSC population which can impact the myeloma progression and development of MBD. Moreover, our preclinical results support the idea that therapeutic targeting of aberrant DNA methylation would result in an anti-myeloma effect and also preserves the appropriate osteogenic differentiation of MSCs to combat myeloma bone disease.

## Acknowledgments

We thank CERCA Programme/Generalitat de Catalunya for institutional support. E.B. was funded by the Spanish Ministry of Economy and Competitiveness (MINECO; grant numbers SAF2014-55942-R and SAF2017-88086-R) and the Multiple Myeloma Research Foundation. M.G. received financial support from the Spanish FIS-ISCIII (PI15/02156) and FEDER. A.G.G is funded by a postdoctoral contract of the Asociación Española Contra el Cáncer (AECC).

## Author contributions

A.G-G., T.L., J.R.-U. and E.B. conceived and designed experiments; A.G-G., T.L., J.R.-U., L.C. and L.S-S. performed experiments; A.G-G, T.L. and F.C.-M. performed biocomputing analysis; M.M., L.S-S., X.M., J.O., C.O-d-S., E.S.J-E., X.A., F.P. and M.G. participated in the data acquisition (performed patient selection, provided drugs, samples, animals and facilities); A.G-G, T.L., J.R-U. and E.B. analyzed and interpreted the data; A.G-G, T.L and E.B. wrote the paper. All authors read and approved the final manuscript.

